# Analysis of Antibody Hybridization and Autofluorescence in Touch Samples by Flow Cytometry: Implications for Front End Separation of Trace Mixture Evidence

**DOI:** 10.1101/045948

**Authors:** M. Katherine Philpott, Cristina E. Stanciu, Ye Jin Kwon, Eduardo Bustamante, Susan Greenspoon, Christopher J. Ehrhardt

## Abstract

The goal of this study was to survey optical and biochemical variation in cell populations deposited onto a surface through touch or contact and identify specific features that may be used to differentially label and then sort cell populations from separate contributors in a trace biological mixture. Cell characterizations initially focused on two different protein systems, Human Leukocyte Antigen (HLA) complex and cytokeratin (CK) filaments. Hybridization experiments using pan and allele-specific HLA antibody probes showed that surface antigens on cells transferred from the palmar surface of volunteers are largely unreactive, suggesting that they cannot be used to differentiate cell populations in a touch mixture. Touch samples were also hybridized with the pan-CK probe AE1, which targets CK proteins 10, 14, 15, 16 and 19. Fluorescence levels of AE1 hybridized cells were observed to vary across donors, although these differences were not consistent across all sampling days. We then investigated variations in red autofluorescence profiles (650-670nm) as a potential signature for distinguishing contributor cell populations. Although distinct differences in red autofluorescence profiles were observed ‐‐ with one donor consistently exhibiting higher levels of fluorescence than others ‐‐ some variation was also observed in touch samples collected from the same individual on different days. While this suggests that contributor touch samples cannot be defined by a discrete level of autofluorescence, this attribute may still be a useful means of isolating contributors to some touch mixtures. To test whether these observed optical differences could potentially be used as the basis for a cell separation workflow, a controlled two person touch mixture was separated into two fractions via Fluorescence Activated Cell Sorting (FACS) using gating criteria based on intensity of 650-670nm emissions, and then subjected to DNA analysis. STR typing of the sorted fractions provided partial profiles that were consistent with separation of individual contributors from the mixture.

## Introduction

Analysis of ‘touch’ or trace epithelial cell mixtures is a significant problem for DNA caseworking units. Currently, interpretation of STR profiles containing multiple contributors requires time-consuming and frequently subjective procedures that often decrease the probative value of the evidence and can lead to its total loss. Although probabilistic genotyping systems can perform analyses on complex mixtures which are superior to human analysis, implementation of these systems poses a number of challenges (e.g. cost; time requirements; legal skirmishes over proprietary software; difficulties associated with communicating probabilistic information to a jury), mis-estimation of the number of contributors to a sample can affect probabilistic results [1], and are limits as to the number of contributors that can be successfully disentangled [2]. There remains a considerable need for front end techniques that can separate cell populations from different contributors prior to DNA analysis thereby facilitating the generation of single source STR profiles and/or simplifying multi-contributor samples, particularly for mixtures of greater than three individuals [3].

Towards this end, a number of methods exist for isolating cells from a mixture including laser capture microdissection, microfluidic manipulation, and fluorescence activated cell sorting [4–7]. These techniques typically take advantage of either morphological or immunochemical variation between cells and have proven to be effective for resolving mixtures with multiple cell types (e.g., blood-saliva [7], sperm-epithelial [6,8]) or mixtures with only one cell type [4]. However, few studies have tested front-end separation strategies on ‘touch’ mixture samples consisting solely or largely of epidermal cells which have vastly different biological and structural properties from other forensically relevant cell types (e.g., vaginal, buccal, white blood cells).

Specifically, sloughed epidermal cells (aka corneocytes) come from the outermost layer of the skin, the stratum corneum. Prior to shedding, corneocytes have undergone a process of terminal differentiation as they migrate to the outer layer whereby they have lost their cellular organelles including their nuclei [9,10]. Additionally, various keratin molecules accumulate both within and upon the surface of the cell as it becomes encased within an extracellular matrix composed of lipids and hydrophobic proteins that contribute to the skin’s barrier function [11,12]. The progressive keratinization of epidermal cells and degradation of their intracellular components may pose considerable obstacles to the development of immunochemical techniques to differentiate contributors in touch mixture samples. Surface antigens, which can be targeted for selective labelling of individual cell populations in a forensic mixture [4,7,8], have variable reactivity in keratinocytes originating from different epidermal layers (e.g., basal layer is more reactive than spinous layer) [13–15], and their utility has yet to be explicitly evaluated in touch samples.

Therefore, the objective of this study was to characterize the optical properties and immunochemistry of cells recovered from touch or contact biological samples with the overarching goal of identifying biomolecular targets that may be used to differentiate, and ultimately separate, epidermal cell populations from different individuals. We initially focused on the reactivity of epidermal cells to antibody probes that target two different protein classes: the Human Leukocyte Antigen (HLA) complex and cytokeratins (CKs). We followed these experiments with a survey of intrinsic fluorescence of epidermal cells from different contributors at red wavelengths (650nm-670nm). Next, in an effort to physically isolate contributor cell populations in a controlled two-person ‘touch’ mixture, we used the observed inter-contributor variation in autofluorescence profiles to develop gating criteria for subsequent fluorescence activated cell sorting (FACS). Finally, we processed the sorted cell populations using forensic DNA analysis methods, and compared the STR profiles of each fraction against profiles from the unsorted mixture and the contributor reference samples in order to evaluate the efficacy of separation.

## Methods

### Sample Collection

Touch samples were obtained pursuant to VCU-IRB approved protocol ID# HM20000454_CR. Volunteers were asked to rub a sterile polypropylene conical tube (P/N 229421; Celltreat Scientific) using their palm and fingers for five minutes. Cells were collected from the surface with sterile pre-wetted swabs (P/N 22037924; Fisher Scientific) followed by dry swabs. A total of six wet swabs and two dry swabs were used to sample the entire tube surface. To elute the cells into solution, the swabs were manually stirred then vortexed for 15 seconds in 10 mL of ultrapure water (18.2 MΩ·cm). The entire solution was then passed through a 100 μm filter mesh prior to AMNIS imaging, antibody hybridization, and/or flow cytometry. Separate aliquots of the resulting cell solution were used for each analysis method, with the exception of the samples used in mixture separation studies which were collected as described below.

For the two-person mixture studies, each donor rubbed a tube as described above. The entire surface area of the tube (excluding the cap) was swabbed with one slightly wetted cotton-tipped swab followed by one dry swab. The swabs were then eluted in 2mL of sterile water,vortexed for 15 seconds, and passed through a 100 μm mesh filter. An 860 uL aliquot of each donor’s touch cell solution was combined to create a 1:1 mixture (by vol.) for flow cytometry analysis, gating, and subsequent sorting via FACS. Another 200 uL from each donor was combined to create a mixture that would proceed directly to DNA analysis without sorting (i.e. to develop an unsorted mixture profile for comparison). The remaining cell solution for each of the two donors was utilized for microscopic imaging studies.

### Cellular Imaging

In order to preliminarily assess the nature of cellular material comprising our touch samples, we analyzed the morphological and optical characteristics of samples from three of our donors. First, an aliquot of each donor’s touch cell solution (~500 μL) was sorted into ‘large cell’ and ‘small cell’ fractions (i.e. ‘K’ and ‘D’ fractions, respectively) using a BD FACSAria™ Ilu (Becton Dickinson) flow cytometer with 488 nm and 633 nm coherent solid state lasers, and set to the following channel voltages: FSC,200V; SSC,475V. Cells and other events were sorted into large and small cell fractions based upon their forward scatter (FSC) and side scatter (SSC) profiles. Sorting was performed until at least 1,000 events (cells) were collected into each fraction.

Each fraction was then analyzed using an Amnis^®^ Imagestream X Mark II (EMD Millipore) equipped with 488nm and 642nm lasers. Images of individual cell events were captured in the Brightfield channel as they passed through this specialized flow cytometer. Additionally, we conducted microscopic surveys of autofluorescence in individual cells by activating both lasers and the APC channel detector (642-745nm). Magnification and focus settings varied with cell size. Cell images were analyzed and exported with the IDEAS^®^ Software (EMD Millipore).

### Antibody Hybridization

Three milliliter aliquots of ten donors’ touch cell solutions were centrifuged at 5,000xg for five minutes. The resulting cell pellets were then redissolved in ~100 μL of supernatant and incubated for 10 minutes with 1 μL of Human Fc Receptor block (Cat# 130-059-901, Miltenyi Biotec) to increase the specificity of antibody binding before reaction with either HLA or CK probes. For HLA hybridizations, cells were incubated with mouse anti-human monoclonal antibody (mAb) HLA-ABC-FITC (Cat# 311403, BioLegend) for 30 minutes. Cells incubated with anti-mouse IgG2a-FITC (Cat# 343303, BioLegend) for 30 minutes served as the isotype control for these experiments. Cells were then washed once in 1x FACS buffer [PBS supplemented with 2% Fetal Bovine Serum (FBS, Cat# 100-106, Gemini BioProducts) and 10% Sodium Azide (Cat# S2002, Sigma-Aldrich)] and re-suspended in the same solution until flow cytometry analysis.

For CK hybridization experiments, cells were incubated with anti-acidic cytokeratin probe (‘AE1’ (recognizes CKs 10, 14, 15, 16 and 19), Cat# 14-9001-80, Affymetrix eBioscience) for 30 minutes followed by reaction with a secondary antibody, anti-mouse IgG1-APC (Cat# 17-4015-80, Affymetrix eBioscience). We used anti-mouse IgG1-APC (Cat#17-4714-42,Affymetrix eBioscience) to create the isotype control for AE1 experiments, incubating for 30 minutes. As before, cells were washed once and then resuspended in 1xFACS buffer prior to analysis.

### Flow Cytometry and Fluorescence Activated Cell Sorting

For HLA and CK studies, flow cytometry analysis was performed on the BD FACSCanto™ II Analyzer (Becton Dickinson) equipped with 488nm and 633nm lasers. Channel voltages were set as follows: Forward Scatter (FSC, 150V), Side Scatter (SSC, 200V), Alexa Fluor 488 (FITC, 335V), Phycoerythrin (PE, 233V; PE-Cy5, 300V; PE-Cy7, 400V), and Allophycocyanin (APC, 250V). For each experiment, 10,000 total events were collected for analysis. Data analysis was performed using FCS Express 4 Flow Research Edition (De Novo Software).

Intrinsic fluorescence studies of six donors’ touch samples and Fluorescence-Activated Cell Sorting (FACS) of two-person epidermal cell mixtures were performed on one of two BD FACSAria™ Ilu (Becton Dickinson) flow cytometers, each employing 488 nm and 633 nm coherent solid state lasers. On each instrument, channel voltages were set as follows: FSC, 200V; SSC, 475V; APC, 400V. Cell events falling into the ‘large cell’ gate (i.e. ‘K’ fraction) were analyzed for red autofluorescence (650-670nm), again using FCS Express 4 Flow Research Edition. For the mixture sample, sorting gates were set to enrich for each of the two contributors in the mixture based on their individual autofluorescence profiles (‘P9’ and ‘P10’ regions of the fluorescence histograms shown in Figure 6). We collected 15,406 events/cells in ‘Sort A’ and 10,607 events/cells in ‘Sort B’.

### DNA Extraction, Purification and Quantitation

Sorted samples were centrifuged at 10,000 xg for 15-20 minutes to pellet cells. The supernatant was concentrated onto a YM-100 Microcon filter (P/N 42413, EMD Millipore) and eluted in 25 μl of sterile distilled water, then re-combined with the cell pellet. These samples, as well as the reference samples (unsorted mixture and donor buccal samples) were each lysed and purified using the DNA IQ System (Cat# DC6701, Promega) following the VA-DFS standard protocols [16]. DNA extracts were quantitated using the Plexor HY System kit (Cat# DC1001, Promega) coupled with the Stratagene MX3005P Quantitative PCR Instrument and Plexor Analysis Software.

### STR Amplification and Profiling

We used the PowerPlex^®^ Fusion System kit (Cat# DC2402, Promega) to amplify STRs in an ABI 9700 thermal cycler, following the manufacturer’s protocols. Capillary electrophoresis was performed on the ABI 3500 xL Genetic Analyzer (Life Technologies) as described in the instruction manual, and resulting data was analyzed using GeneMapper ID^®^ ‐X v1.4 Software (Life Technologies) according to the manufacturer’s recommendations. The analytical thresholds used to interpret the resulting data were dye specific and set at 88 relative fluorescent units (RFU) for fluorescein, 74 for JOE, 114 for the TMR-ET, and 80 for CXR-ET. The stochastic threshold was set at 396 RFU.

## Results

### Morphological Characterizations

Initial optical characterizations of cell solutions recovered from touch samples showed two distinct populations. Individual events within the ‘K’ gate measured ~20-40 μm in diameter. The size and morphology of the subset of large cell (K) fraction events imaged with AMNIS (typically several hundred cells per sample) appeared to be consistent with intact keratinocytes (top two rows of cell images in Figure 1); we did not observe any cells with features that would suggest the presence of other epithelial cell types (e.g., buccal). Evidence of folded or rolled cells was also observed in the K population which likely reflects physical deformation of some cells during surface swabbing. Events within the ‘D’ population were typically less than 10 μm. Their size and overall variable morphology in AMNIS images (bottom two rows in Figure 1) suggest that these events represent cell fragments, biological debris, or non-cellular particles such as hairs or fibers. The distribution of cell events within the K-population vs. D-population were observed to show considerable variation both between donors and between sample replicates from the same individual, consistent with previously published data for these samples [17].

**Figure 1.**
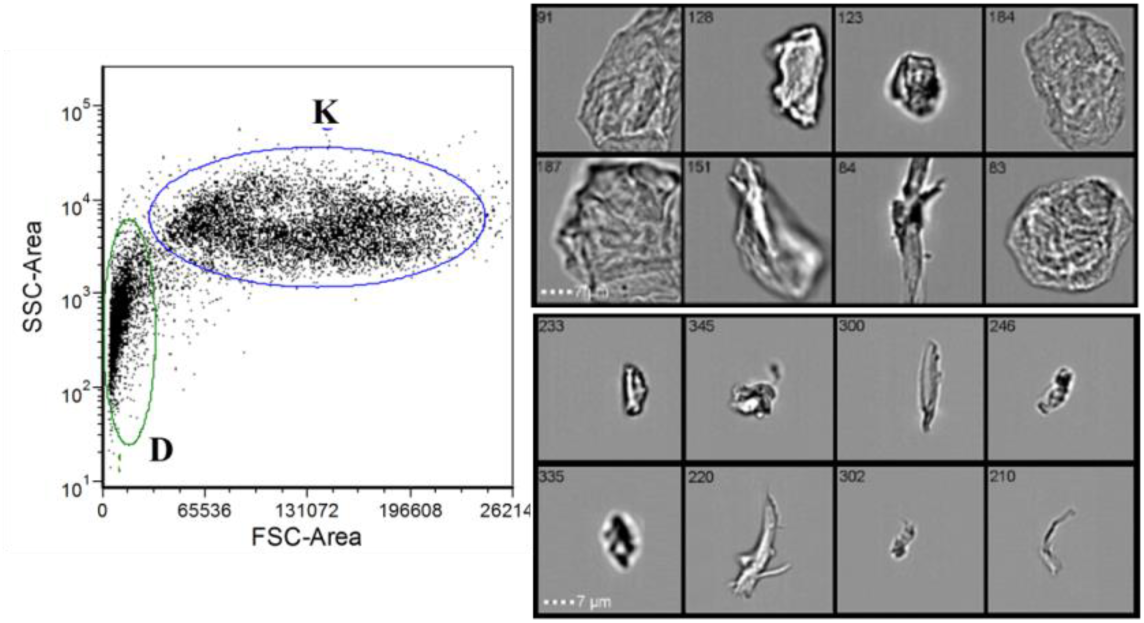
Optical characterization of a touch cell solution. Forward scatter and side scatter plot of all cell events showing ‘K’ and ‘D’ subpopulations (left). Images of individual cell events using AMNIS instrumentation (right). The top two rows are sampled from the K subpopulation and the bottom two rows are from the ‘D’ subpopulation.

### Antibody Labeling Experiments

Touch samples from ten donors were each hybridized to a fluorescently-labelled pan-HLA antibody that recognized all antigens within the A, B, and C protein classes. Probe-hybridized cells displayed no increase in average fluorescence when compared to unlabeled cells or isotype controls (Figures 2a–c). Similar results were obtained when HLA probes specific for the A*02 allele were hybridized against cells that screened positive for the A*02 allele (data not shown). Although there was no obvious change in fluorescence after probe hybridization, some differences in the distribution of FITC channel intrinsic fluorescence values were consistently observed from one donor to the next (e.g., compare purple and dark blue histograms in Figures 2a–c).

**Figure 2.**
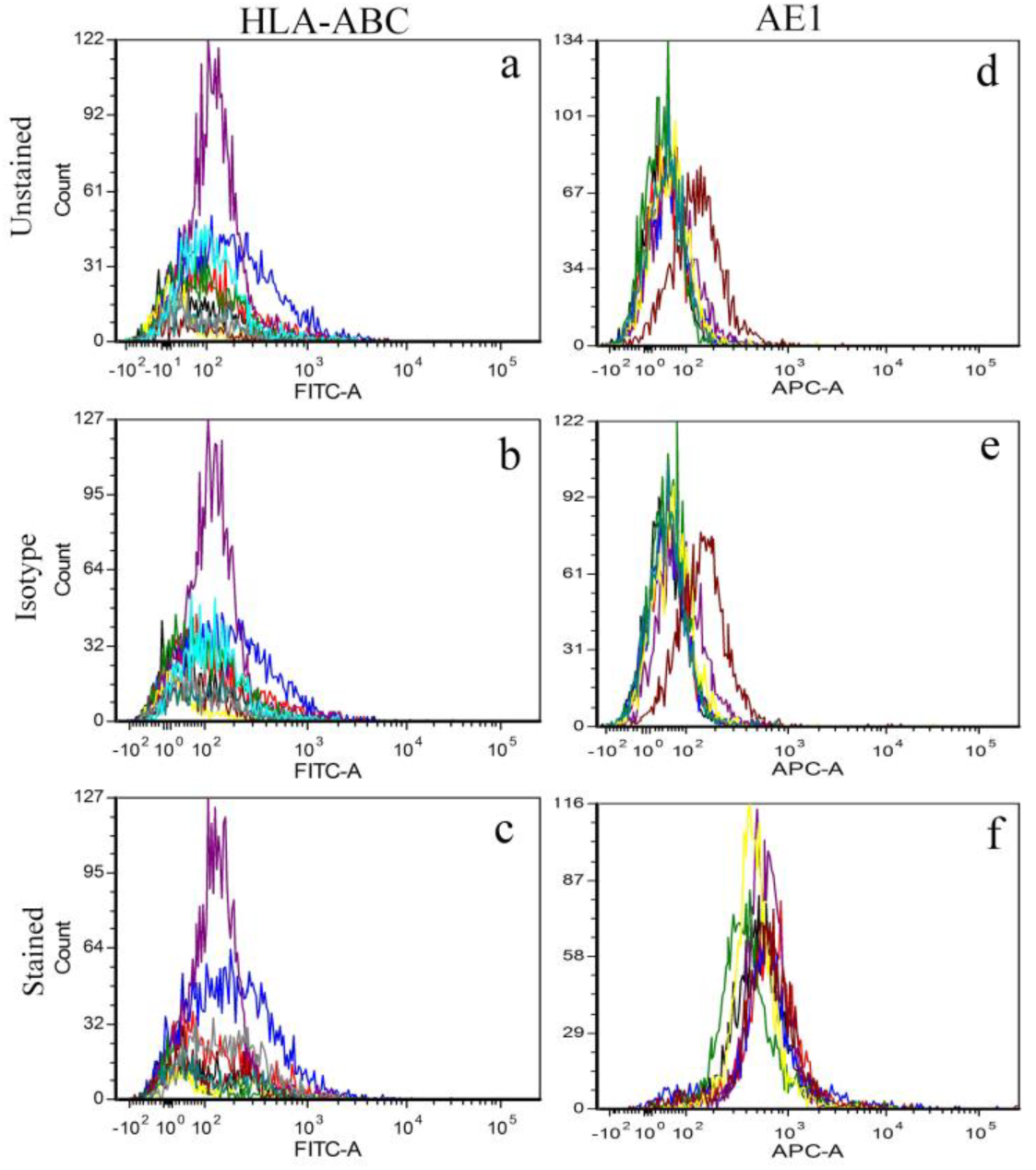
Hybridization of touch samples with HLA and CK antibody probes. Few differences were observed between samples hybridized with pan-HLA probe and unstained samples/isotype controls, indicating that the touch samples failed to uptake the probe (panels a-c). In contrast, all touch samples exhibited uptake of AE1 cytokeratin antibody probe, with slight differences observed in binding efficiency across contributor cell populations (panels d-f).

In contrast, experiments using AE1 cytokeratin antibodies show probe uptake for each of ten donor samples tested when compared against unstained cells and isotype controls (compare Fig 2f to Figs. 2d and 2e). We observed slight inter-individual variation in binding efficiency. Mean fluorescence intensities ranged between 417 and 663 relative fluorescence units (RFUs), with all donors exhibiting significant overlap in their histogram profiles. Of note, one donor cell population showed higher average levels of intrinsic APC channel fluorescence compared to other donors (maroon histogram in Figs. 2d–e). Interestingly, this donor cell population displayed one of the lowest probe binding efficiencies of those surveyed (maroon histogram in Fig. 2f). When touch samples from a subset of these donors were monitored for changes in AE1 uptake between sampling days, we found that the efficiency of probe binding varied from day to day, with samples from one donor in particular exhibiting discernibly higher fluorescence than other donors on two of the four collection days (Figure S1, red histograms).

### Intrinsic Fluorescence Surveys

Next, we examined variation in intrinsic fluorescence at red wavelengths (~650-670 nm) as a potentially discriminating characteristic for cell populations from different individuals. This wavelength was chosen based on initial observations in the course of antibody hybridization studies that unstained cell samples from some contributors showed higher mean fluorescence intensities than others (Figs. 2d and 2e (maroon histograms); Figure S2 [18], and data provided in [19]). Further, images collected during microscopic surveys indicated a number of cells exhibiting APC channel fluorescence (Fig. 3); among the images captured, we observed some noticeable differences in fluorescence intensities between cells from different individuals (e.g., Fig. 3, comparison between E15 (particularly frames 13 and 20) and D02 cell events.

**Figure 3.**
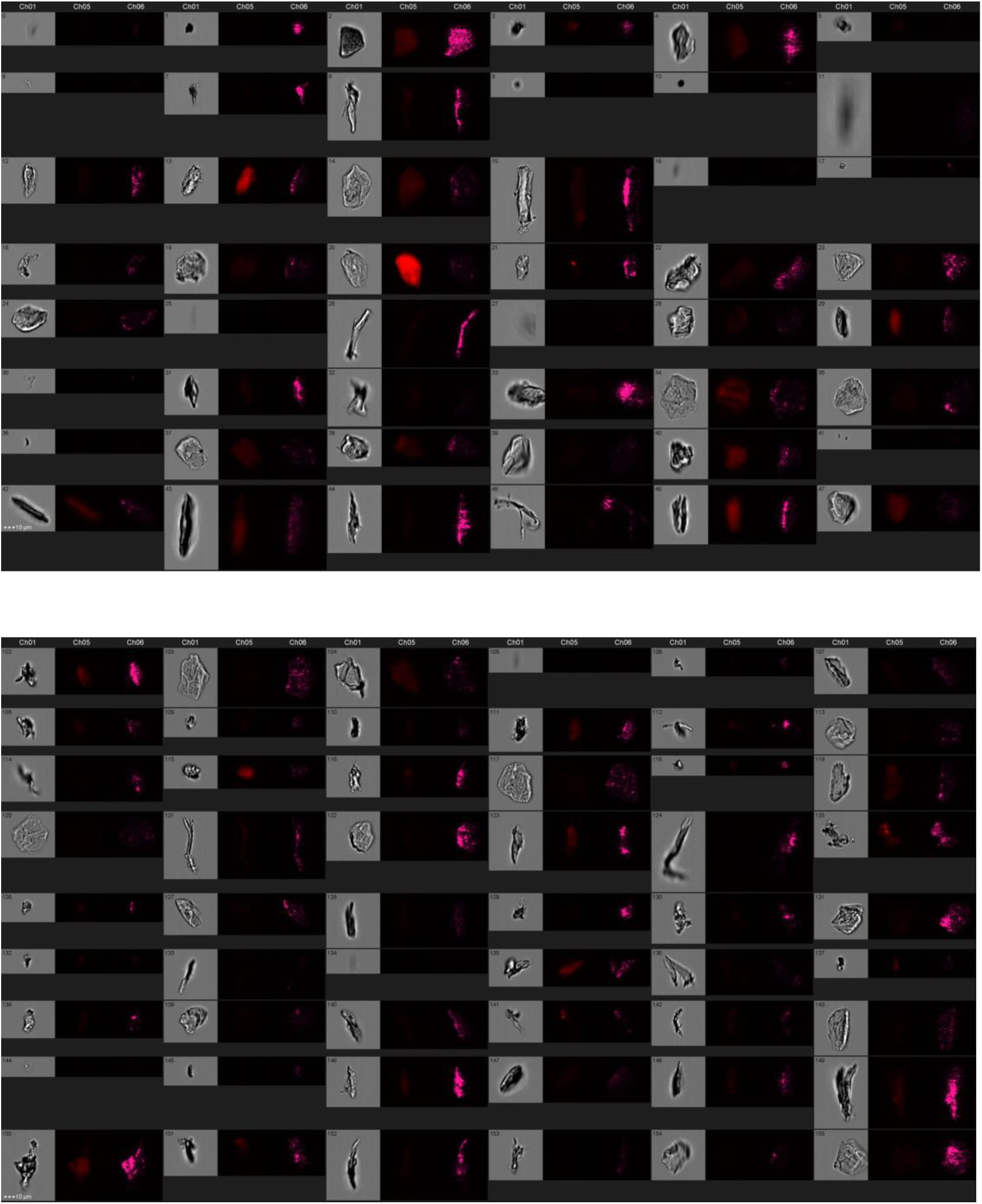
AMNIS imaging of individual flow cytometry events from the large (“K”) fraction of touch samples from two different contributors, E15 (top) and D02 (bottom). Each event was visualized in three different microscopic settings: Brightfield (left image in gray), APC channel fluorescence (middle image shown in red), and side scatter (right image shown in purple).

Intrinsic fluorescence profiles (APC channel) were developed from six donors’ touch samples, with different subsets of these individuals sampled and analyzed on three different days; results are shown overlayed and grouped by sampling day in Figure 4. Significant overlap was observed between many of the donors on each sampling day. However, touch samples from one contributor, E15 (red histogram in each panel), consistently contained a number of cells with higher fluorescence intensity than cells from other contributors. Microscopic surveys of individual cell events from contributor E15 showed red autofluorescence associated with what appear to be intact corneocytes (Figure 3). Fluorescence was also observed associated with other flow cytometry events, which could be rolled or fragmented cells, or possibly non-cellular material such as fibers.

**Figure 4.**
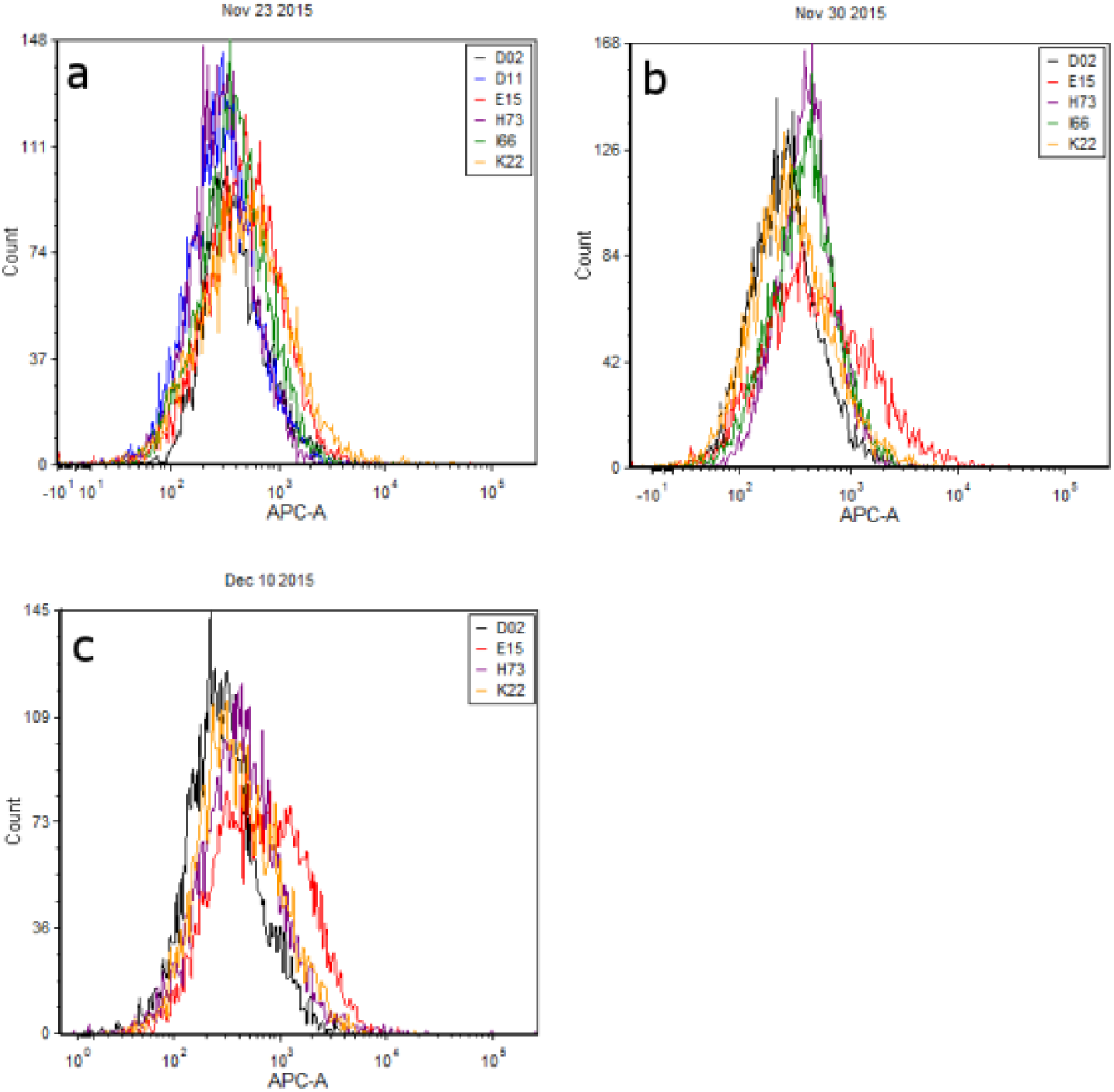
Overlayed red fluorescence (650-670nm) histograms for cell populations from touch samples. Each panel (a-c) shows a different combination of donor cell populations sampled and analyzed on the same day.

To investigate the consistency of autofluorescence signatures, touch samples were collected from donors E15 and D02 on seven additional days and analyzed for red autofluorescence. Results showed that the degree of differentiation (or conversely overlap) between autofluorescence profiles varied considerably across days (Figure 5). Nonetheless, the mean APC channel fluorescence of E15 cell populations was consistently higher than D02 populations.

**Figure 5.**
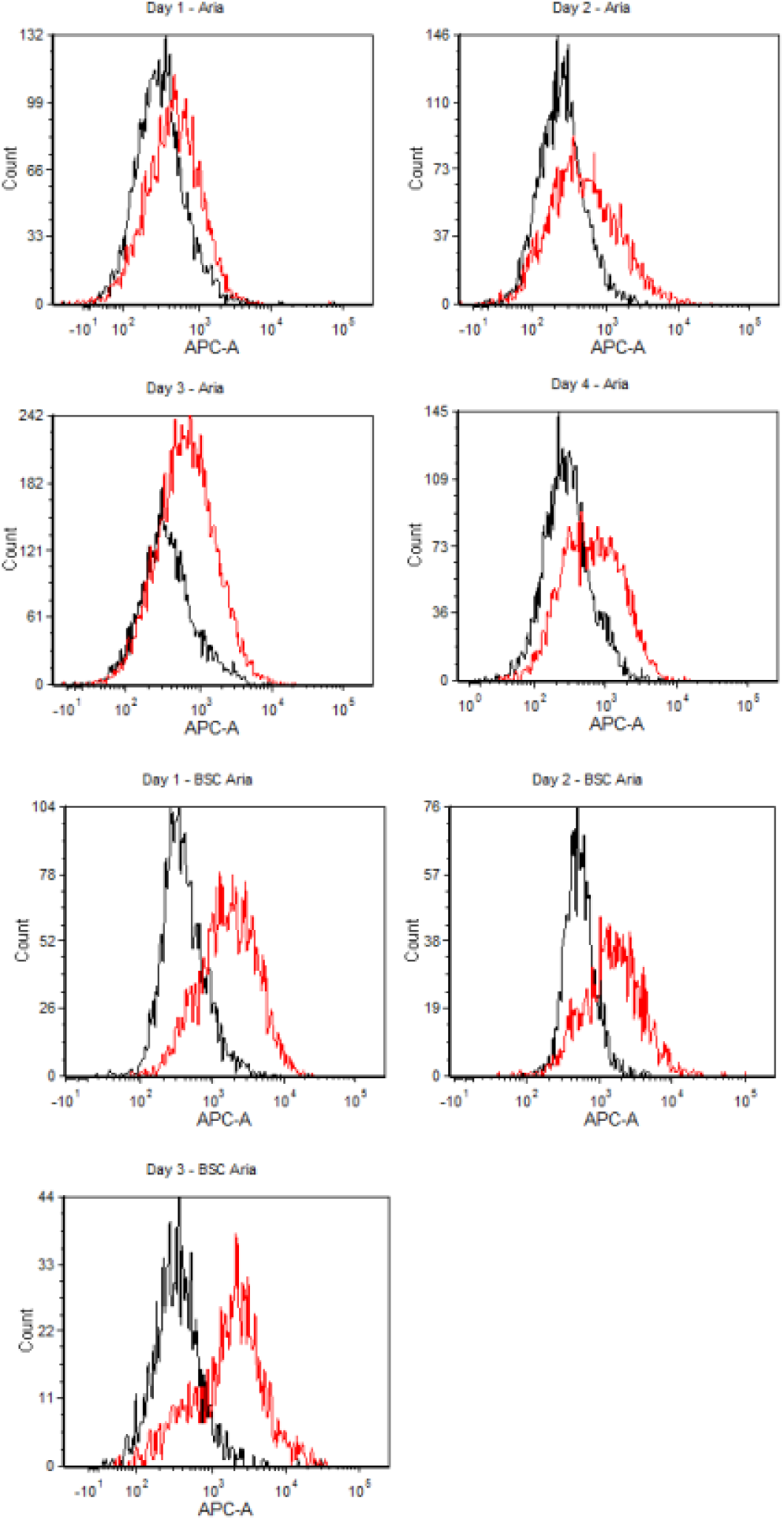
Overlayed red fluorescence histograms for two contributors, D02 (black) and E15 (red), across seven independent sampling days.

### Two-person mixture study

Mixtures of cells deposited by donors D02 and E15 were sorted into separate fractions via FACS according to the gating criteria shown in Fig. 6, and then subjected to DNA analysis. “Sort A” and “Sort B” – the cell fractions that met gating criteria derived from intrinsic fluorescence measurements of cells from donors D02 and E15 (respectively) – each produced a partial profile (Table 1). The high degree of dropout, and possible drop-in alleles observed are consistent with extremely low level of template DNA detected in each cell fraction (<50pg).

**Figure 6.**
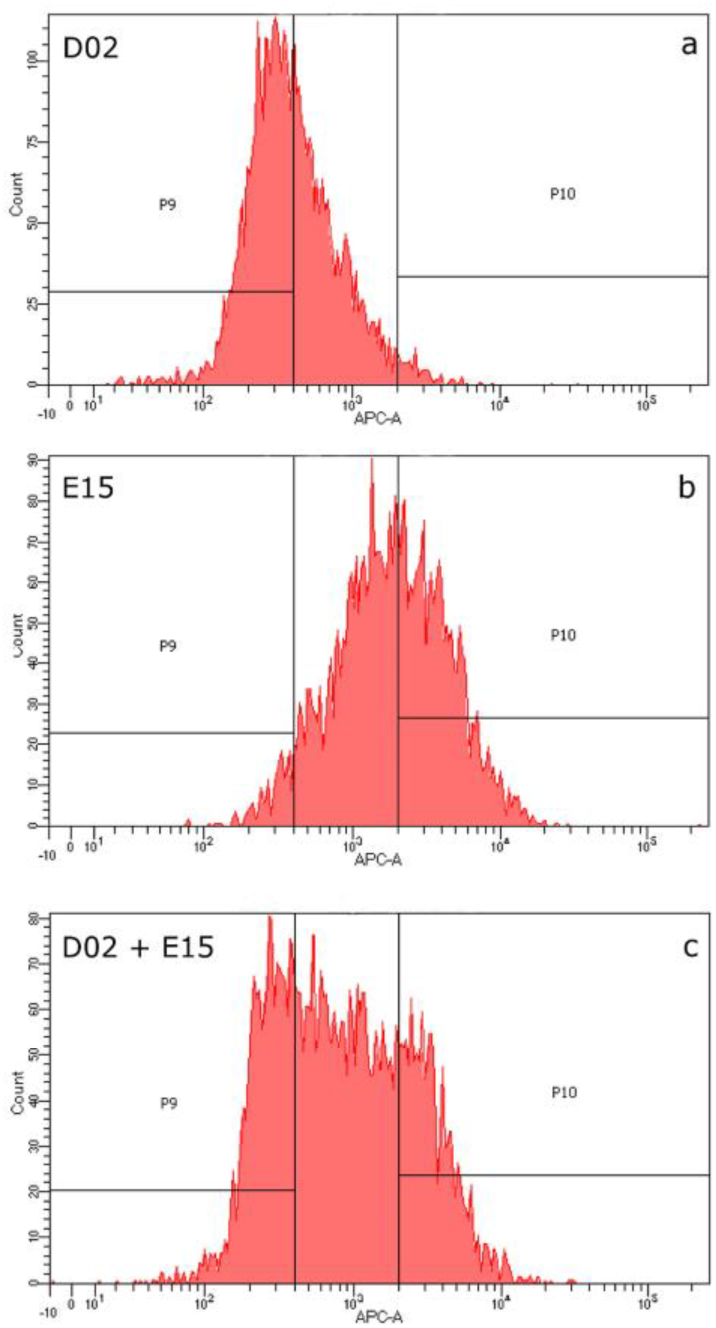
Sorting gates used for FACS based on APC channel intrinsic fluorescence. Histogram profiles for single source samples (panels a, b) were used to define two sorting gates, P9 and P10. These gates were positioned such that cell populations from D02 and E15 would be enriched relative to each other in the two cell fractions. Panel c shows the sorting gates plotted against the histogram profile of the two-person cell mixture prior to sorting.

**Table 1.**
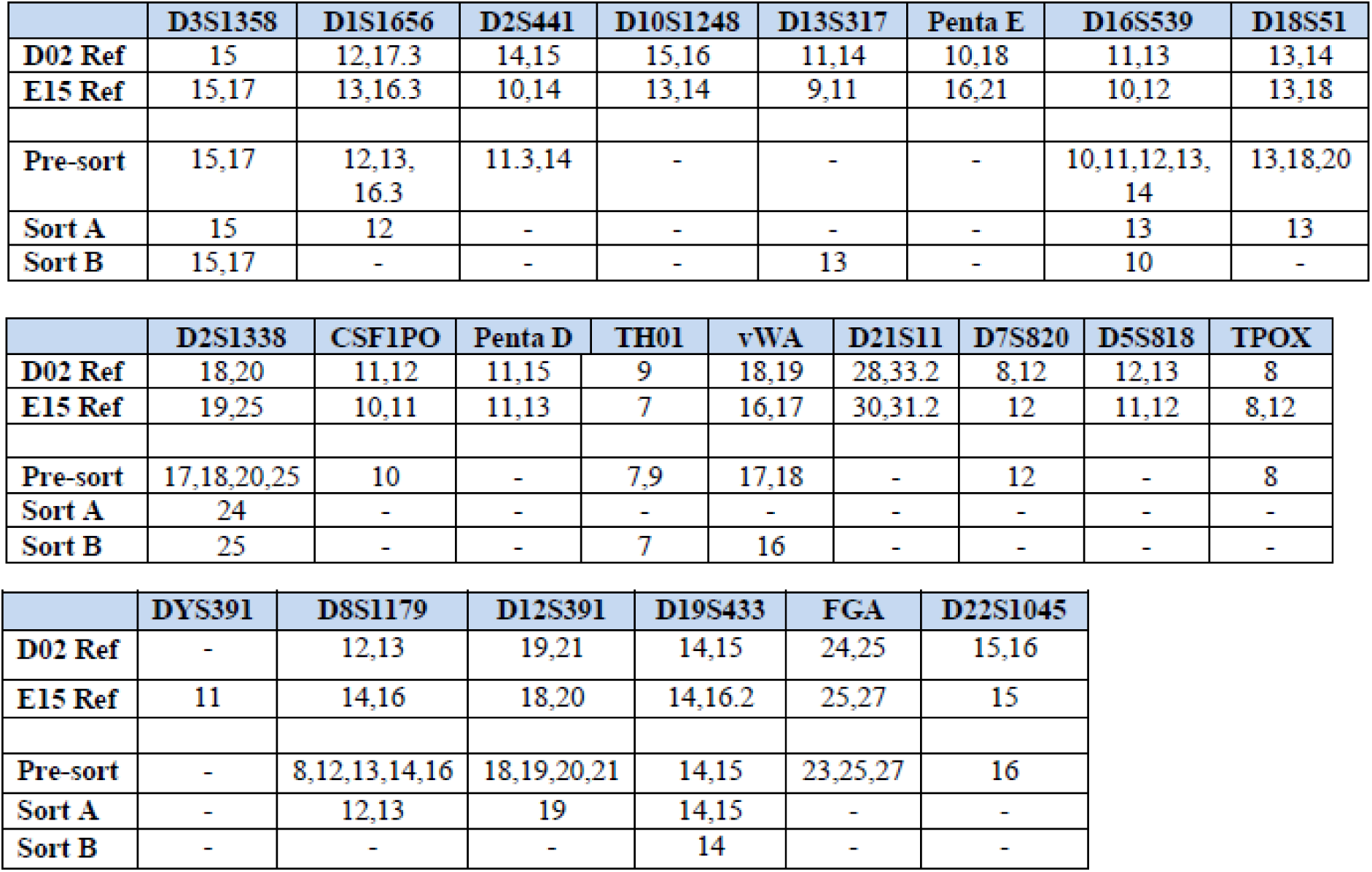
Powerplex fusion profiles developed from donors D02 and E15 reference samples (buccal), unsorted mixture of cells deposited by D02 and E15, and sorted fractions. “Sort A” is the cell fraction that met the gating criteria based upon D02’s intrinsic fluorescence profile, and “Sort B” is the cell fraction that met the gating criteria based upon E15’s intrinsic fluorescence.

All alleles detected in Sort A were consistent with donor D02 with the exception of a single 24 allele at locus D2S1338, which did not originate from donor E15 and is likely a drop-in allele; none of E15’s obligate (i.e. unique) alleles were detected in the DNA profile developed from Sort A. Likewise, all alleles detected in Sort B were consistent with donor E15 with the exception of a single 13 allele at locus D13S317, which did not originate from donor D02 and is likely a drop-in allele; none of D02’s obligate alleles were detected in the DNA profile developed from Sort B.

## Discussion

The objective of this study was to characterize the optical and biochemical properties of touch epidermal cell samples and investigate different cellular properties that may be used to differentiate contributor cell populations in a touch mixture. Flow cytometry data showed that biological material recovered from standard sampling swabs and eluted in solution was composed of intact cells consistent in size with corneocytes (20-40μm) and smaller, irregular events. Single cell imaging of the latter fraction suggests that it is composed of cellular debris, deformed/damaged cells, and fiber fragments that may originate from the collection swab or are associated with the sampled substrate. Over the course of this study we observed both inter‐ and intra-contributor variation in the number of corneocytes detected and their percentage relative to the total number of events in a touch sample, consistent with previous reports [17]. Nonetheless, cell yield was not an issue as touch swabs routinely provided more than 10,000 cells for analysis.

As with our previous studies of controlled touch samples, evidence of other epithelial cell types was not observed or detected (e.g., buccal cells which generally appear larger than corneocytes [20]), although we note that damaged or fragmented cells from other tissues may be difficult to detect with these techniques. Since cell source information can be probative in some cases (e.g. to support or refute allegations of oral contact vs. touching), future research should focus on methods of differentiating and identifying different epithelial cell types. As discussed further below, the same classes of proteins that we surveyed in these studies could potentially be used for this purpose [21], and possibly integrated with other cell targets and/or properties into the kind of flow/FACS methodology that we investigate here, thus permitting simultaneous discrimination between cell types and contributors to a mixture.

We hybridized epidermal cells against two different classes of antibody probe in order to assess whether the target proteins’ variable expression had the potential to differentiate donor cell populations in a touch mixture. Hybridization experiments targeting HLA antigens on the cell surface showed little to no binding to either allele-specific or class-level antibody probes, suggesting that HLA antigens were either not present or were unreactive (Figure 2). The absence of HLA probe interactions in this study is further evidence that the overwhelming majority of cells in these touch samples are fully differentiated keratinocytes, which have been shown to display limited reactivity to HLA Class I probes in contrast to cells derived from deeper layers of the epidermis [13,15] or non-epidermal epithelial cell sources [22].

Of course, there is no such thing as a representative touch sample, and likely some touch samples encountered in casework will include non-corneocyte components such as buccal cells [23], which may prove reactive to cell surface probes. However, we did not detect any such cells in this study or in our previous work [20], which may be characteristic of many of the touch samples recovered in case work. Regardless, before abandoning cell surface antigen targets such as these in touch samples, it may be worth exploring techniques such as preliminary trypsinization to increase immunoreactivity of corneocytes [13,15].

For purposes of the current studies, though, we moved on from HLA probes to test an antibody probe system that targets cytokeratins, which are an important structural component of both differentiating and fully differentiated epidermal cells [12]. Specifically, we utilized AE1 probe which binds to cytokeratin proteins 10, 14, 15, 16, and 19. We found that touch samples consistently hybridized to the AE1 probe, albeit donors displayed slight variation in binding affinity (Figs. 2, S1). Across sampling days, the degree of variation occasionally increased (Figs. S1a and S1c), however, we observed that the difference was sometimes minimal (Figs. 2, S1b and S1d). These results suggest that cytokeratin expression – at least on the pan-level that is capable of being explored with a probe such as AE1 – may not present a consistently useful means of discriminating between individuals.

However, individual CK probes may prove more discriminating than pan probes, e.g., certain cytokeratins are upregulated, and others downregulated, with age [24]. By targeting cytokeratins on a pan level, these differences may be cancelled out. Expression of individual CK proteins has also been used to distinguish between epithelial cell sources (mucosal epithelial cells (buccal or vaginal) from epidermal cells) [21], and could possibly be used in conjunction with flow cytometry to detect the presence of, and potentially isolate, non-epidermal cell types in touch samples (or, background levels of skin cells in a non-touch sample). Future efforts should continue to investigate intrinsic and/or environmental factors that can contribute to differences in cytokeratin expression (e.g., [25]), as well as whether shifts in CK expression as a function of age, cell source, and other factors can be detected and thus used (individually or in combination) to distinguish cell contributors in touch samples.

Our observation in the course of antibody hybridization studies that intrinsic fluorescence – particularly at red wavelengths – varied between donors led us to pursue this feature for its potential in discriminating between cell populations. In previous studies of autofluorescence in eight donors (three of whom – D02, D11 and I66 – were a part of the current set of experiments), clear differences were observed between donor fluorescence profiles, such that fluorescence-based sorting gates could be conceived that would isolate cells from one or more contributors, to the exclusion or minimal contribution of cells from others [19]. In the current experiments, less distinction was observed between donors (Fig. 4), and it is unclear whether or how much of this may be attributable to differences in instrumentation (the earlier studies utilized the BD Canto platform while the current studies utilized two different FACS Aria instruments; further, the voltage settings for the two sets of experiments differed), the specific donors tested (e.g. J16, the donor who exhibited the highest levels of red fluorescence in the earlier studies, was not available during for this study), or possibly a combination of these factors. The influence and potential impact of day-to-day sample variation cannot be discounted, particularly where previous studies also found fluctuation in fluorescence measurements for a donor whose cells were sampled on multiple days and analyzed on a single instrument (Figure 1f in [19]).

Regardless, in the current study, one donor in particular consistently exhibited higher red fluorescence than other donors (E15 histograms in Figs. 4 and 5). On some days, this donor’s cell populations exhibited autofluorescence several magnitudes greater than other days: across seven sampling days, median autofluorescence for E15’s cell population ranged from ~500 to 3000 RFUs. On the days when E15’s touch samples emitted the highest red fluorescence, the degree of differentiation from other donors’ cell populations, in particular D02, was the greatest (Fig. 5).

Understanding the factors, both intrinsic and extrinsic to the cell, which may cause shifts in autofluorescence will be an important area of future research. As discussed previously [19], there are a number of endogenous molecules within the stratum corneum that can contribute to autofluorescence [26], including molecules such as porphyrins which have emission maxima similar to what was observed in this study [27,28]. Although microscopic surveys are consistent with some portion of the red autofluorescence signal being associated with apparent corneocytes (Figure 3), we also noted that other, likely non-cellular, fluorescent particles could be found in these samples and may contribute to the overall optical profiles. These included particles consistent with hairs or fibers that were recovered from the K-population of multiple donors. There is also the possibility that other types of exogenous fluorescent compounds (e.g., plasticides [29], chlorophyll [30], or inorganic molecules) could associate with cellular material transferred from the palms and contribute to its autofluorescence properties.

Ultimately, the observed degrees of inter‐ and intra-individual variation in red autofluorescence profiles indicate that this may not be highly discriminating attribute, but do not necessarily negate the utility of this signature in simplifying touch biological mixtures for downstream DNA analysis and interpretation [19]. Depending on the contributors to a given mixture, it may be possible to isolate one or more on the basis of autofluorescence, or to separate a mixture of three or more contributors into two or more simpler mixtures. This raises the possibility that flow cytometry – which is inherently non-destructive – could be used to screen touch mixtures for their susceptibility to be separated into individual components (or at least broken down into less complex mixtures) based on this characteristic. For example, a mixture sample that exhibits two or more peaks (or even a plateau as shown in Figure 6) on a fluorescence histogram would be a more promising candidate for cell separation than one that exhibits a single distinct peak.

However, even a touch sample composed of readily-distinguished cell populations will not necessarily separate cleanly, or produce worthwhile STR data. Our group and others have reported on the characteristically low levels of intracellular genomic DNA recovered from cells deposited on touch surfaces [20,31], which is expected given that keratinocyte differentiation involves programmed breakdown of nuclear DNA prior to cell shedding from the stratum corneum [10,32]. This could pose a challenge for the application of cell-based separation techniques on touch samples. With that in mind, we utilized autofluorescent signatures to sort a controlled touch mixture of donors D02 and E15 via FACS and attempted DNA analysis of the resultant fractions using a standard forensic workflow.

Our preliminary efforts resulted in a partial STR profile for each sorted touch fraction that is (with the exception of a single extraneous allele) consistent with the respective known contributor, indicating that separation of cell populations from the two known contributors on the basis of red autofluorescence was successful. However, the single stray allele in each sort suggests that a very low level of DNA from a third party may have ended up in these fractions. Given the low levels of target template, it is possible that these are examples of allelic drop in during amplification; negative controls were clean but this does not exclude the possibility of this phenomenon. Interestingly, six extraneous alleles (i.e. not from D02 or E15) were detected in the reference (unsorted) mixture (Table 1). None of these alleles showed up in profiles developed from Sort A or Sort B. These could be instances of drop in (11.3 at D2S441, 20 at D18S51 and 8 at D8S1179) and pronounced stutter (17 at D2S1338, 14 at D16S539, and 23 at FGA) resulting from low levels of DNA template in the touch mixture. It is also possible that these alleles are derived from extracellular DNA (which would not be expected to show up in sorted fractions) that was transferred to the palms of D02 or E15 before they deposited their touch samples, particularly in light of studies demonstrating the prevalence of extracellular DNA in touch samples [20,31,33].

The high degree of allelic dropout observed in the sorted fractions is not unexpected given the nature of the biological material being analyzed – shed epidermal cells. However, there are several areas in our methodology where adjustments could be made to improve DNA yield and/or maximize the use of the DNA that is present, and thus produce more complete DNA profiles from sorted fractions. For example, we utilized a standard forensic DNA analysis protocol on sorted samples, which could be modified in various ways to increase efficiency (e.g. by reducing extract volume and/or concentrating post quantitation). Moreover, these controlled touch mixtures were split into aliquots to be used for differing purposes during these exploratory studies (e.g. microscopic imaging, FACS, DNA analysis without sorting). As such, only a fraction of the cells collected from touched surfaces were submitted to FACS; if more (or all) of the touch samples were utilized for this purpose, each fraction would likely contain more cells for downstream STR profiling.

Further, by designing the sorting gates in this study with an eye toward producing single source profiles, we sacrificed maximal cell recovery for purity of the sort. As can be seen from Figure 6, gate P9 was designed to capture D02’s cells while excluding most of E15’s cells, and gate P10 was designed to capture E15’s cells while excluding most of D02’s. However, approximately half of each of D02 and E15’s cells went unsorted in the middle area between the two gates. With touch samples, and the associated difficulties related to intracellular DNA yield from corneocytes, it may make sense to shift the gating calculus we used for other types of biological material [4]. Instead of designing gates to produce single source profiles, one might strike a balance between cell recovery and production of simple mixtures with easily discernable major components.

For example, if the gates in Figure 6 were set so that all cells in the D02-E15 mixture fluorescing less than 1000 RFU were sorted into Sort A, and those fluorescing at or greater than 1000 RFU were sorted into Sort B, this should result in recovery of all cells from the mixture between the two fractions. Note that while most of D02’s cells exhibit fluorescence below 1000 RFUs, a few cells fluoresce at a higher intensity (Fig 6a); conversely, while most of E15’s cells exhibit fluorescence above 1000 RFUs, a few cells fluoresce at a lower intensity (Fig 6b). Thus, while each fraction sorted in this manner will contain some cells from the untargeted contributor, resulting in a mixture, the major contributor should be distinguishable and consistent with the vast majority of cells in the simplified mixture created by the sort (D02 in Sort A; E15 in Sort B).

One of the biggest drivers of cell loss in our methodology may be the retention of cellular material in the collection swabs following manually stirring and vortexing in water to elute the cells into solution. The challenge of maximizing DNA yield from collection swabs has been explored by a number of researchers in the forensic sciences, though many of the protocols are not applicable where, as here, cells need to remain intact during elution [34]. Future work should continue to test different elution protocols to maximize cell recovery; optimized buffers [35] and the incorporation of enzymes such as cellulase to break down cotton and encourage the release of cells [36] may hold promise. To the extent that some number of cells will undoubtedly remain trapped despite methodological adjustments, subsequent studies should investigate whether and how information derived from this biological material may be exploited. At very least, this unsorted mixture data may be used to give context to STR profiles developed from sorted cell fractions; in some cases, the combination of sorted and unsorted DNA data may increase the overall probative value of a sample.

Finally, because a significant portion of the genetic material in many touch samples may be unavoidably extracellular, characterizing the chemical and physical relationship between cell-free DNA and the surface of intact epidermal cells may be an important area of future research.If extracellular DNA associates with epidermal cells, as it has been observed to do in other cell types (e.g., [37]), flow cytometry protocols could potentially be optimized to maintain surface-bound DNA through the cell sorting process. If it emerges that extracellular DNA is not bound to epidermal cells at the time of transfer, this DNA source can be separately collected for typing [20].

## Conclusions

This investigative study marks a starting point for ongoing research into methods that facilitate the separation of touch samples into individual contributor cell populations for downstream DNA analysis. We continue to explore different properties of corneocytes with the goal of identifying a highly discriminating cellular signature (or combination of signatures). While additional research is needed before FACS can be imported as a front end technique in forensic DNA casework, our preliminary results indicate that there are attributes of fully differentiated keratinocytes that can be harnessed to distinguish cell populations from some individuals. A benefit of a feature such as red autofluorescence is that it can be measured without the need for antibody probes or other special reagents, allowing for touch samples to be pre-screened for this trait.

We are also working to optimize our processing methods to maximize both cell yield and DNA yield from sorted cell populations. However, the recovery of even partial profiles from sorted cell solutions may have the potential to enhance the overall probative value of DNA evidence, particularly when analyzed in conjunction with complex mixture data derived from the same sample (e.g. if it is combined with profiles generated from the extracellular fraction and/or cells retained in swabs). Sorted profiles, even if too incomplete to stand alone, may be able to buttress probabilistic claims about the mixture. At very least, this data could provide important investigatory leads, providing clues as to allelic pairings in an otherwise indistinguishable mixture, and potentially narrowing the pool of suspects.

## Acknowledgements

The authors gratefully acknowledge Daniel Conrad and Julie Farnsworth for providing technical assistance for this project.

## Grant information

This project was funded by the National Institute of Justice Award number 2013-DN-BX-K033 (PI: Ehrhardt). Flow cytometry services in support of the project were provided by the VCU Massey Cancer Center, supported in part with funding from NIH-NCI P30CA016059.

**Figure S1.**
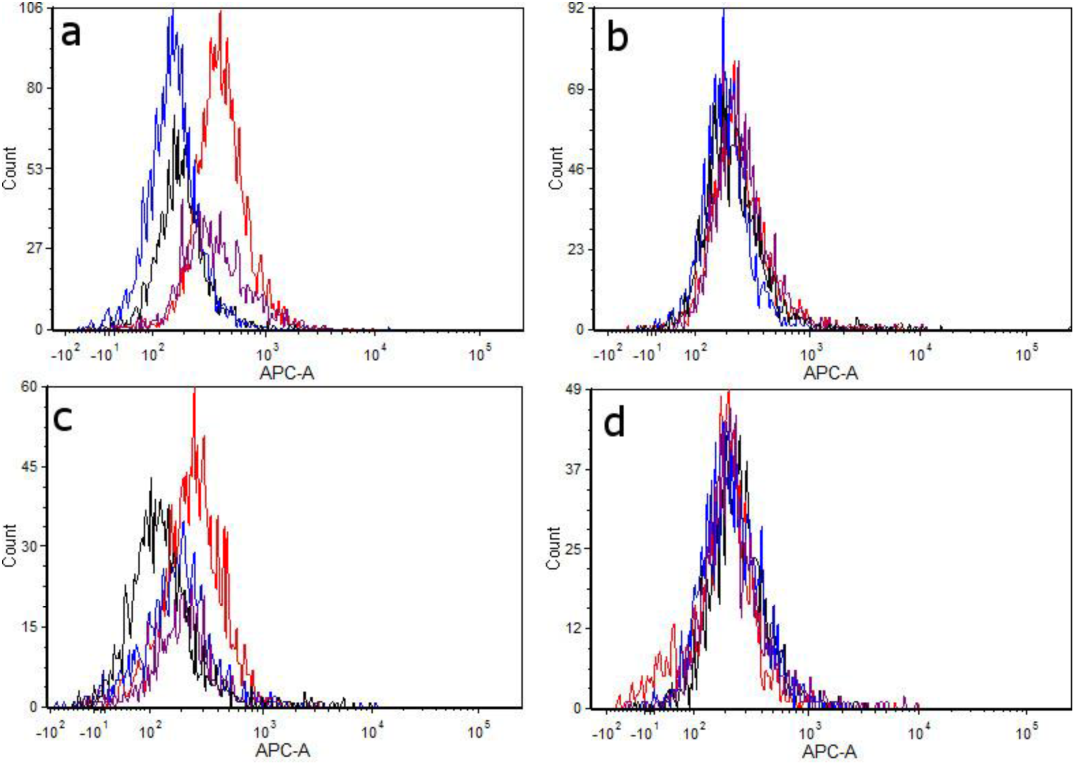
Replicate hybridization experiments using AE1 antibody probe. Touch samples were collected from the same four donors on four different days. On two of the days, differences were observed in the fluorescence profiles exhibited by from cell populations from was observed between donors (a, c). The same differences were not observed for two additional replicate experiments (b, d). Each of the four histogram colors is assigned to a separate contributor cell sample. The same four contributors were examined in each experiment.

**Figure S2.**
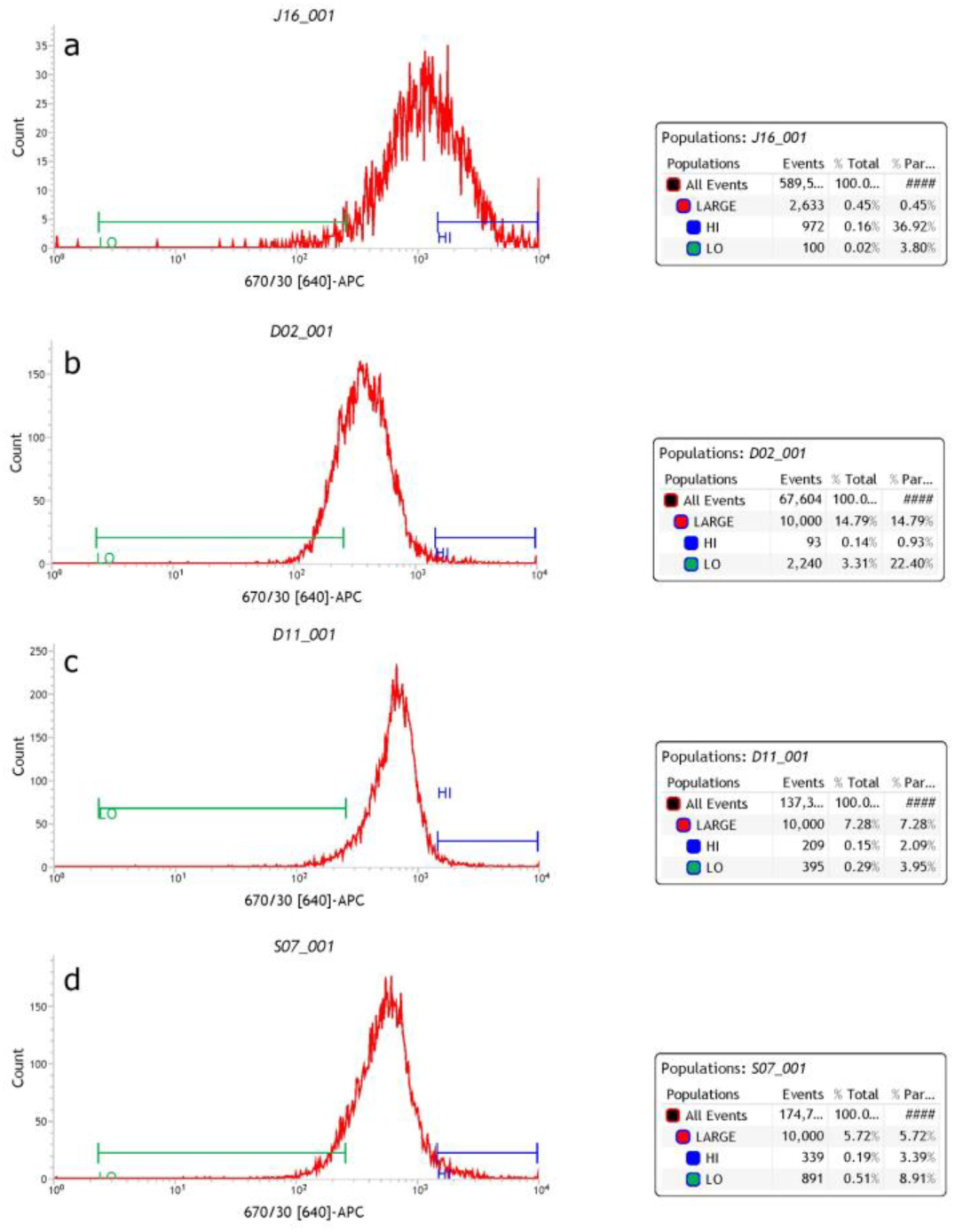
Red Autofluorescence profiles for four donors analyzed using a BD Influx Cytometer. Full source data and method descriptions are given in [18].

